# Identification of H3N2 NA and PB1-F2 genetic variants and their association with disease symptoms during the 2014-15 influenza season

**DOI:** 10.1101/2020.02.20.956979

**Authors:** Deena R. Blumenkrantz, Thomas Mehoke, Kathryn Shaw-Saliba, Harrison Powell, Nicholas Wohlgemuth, Hsuan Liu, Elizabeth Macias, Jared Evans, Mitra Lewis, Rebecca Medina, Justin Hardick, Lauren M. Sauer, Andrea Dugas, Anna DuVal, Andrew P Lane, Charlotte Gaydos, Richard Rothman, Peter Thielen, Andrew Pekosz

## Abstract

The 2014-15 influenza season saw the emergence of an H3N2 antigenic drift variant that formed the 3C.2a HA clade. Whole viral genomes were sequenced from nasopharyngeal swabs of 94 patients with confirmed influenza A virus infection and primary human nasal epithelial cell cultures used to efficiently isolate H3N2 viruses. The isolates were classified by HA clade and the presence of a new set of co-selected mutations in NA (a glycosylation site, NAg+) and PB1-F2 (H75P). The NA and PB1-F2 mutations were present in a subset of clade 3C.2a viruses (NAg+F2P) which dominated during the subsequent influenza seasons. In human nasal epithelial cell cultures, a virus with the novel NAg+F2P genotype replicated less well compared to a virus with the parental genotype. Retrospective analyses of clinical data showed that NAg+F2P genotype viruses were associated with increased cough and shortness of breath in infected patients.

## Introduction

Human influenza epidemics are largely influenced by the emergence and circulation of seasonal viral variants containing mutations that evade pre-existing population immunity (Bedford et al., 2015). This necessitates significant virus surveillance and vaccine efficacy monitoring to guide key public health decision-making and ensure proper preparedness for seasonal epidemics and more infrequent pandemics (Bedford et al., 2015; Flannery et al., 2016; Huzly et al., 2016). The seasonal H3N2 virus variants that became dominant during the 2014-2015 influenza season were associated with an increase in outpatient influenza-like illness hospital visits, hospitalizations, and deaths attributable to pneumonia and influenza (Appiah et al., 2015; Epperson et al., 2014). Increased disease severity was also detected in other countries across the northern hemisphere (Broberg et al., 2015; Donadeu et al., 2015). The increased number of cases correlated with a marked diminution of serum hemagglutination inhibition (HI) and neutralizing (NT) antibodies against seasonal virus isolates (Flannery et al., 2016; Skowronski et al., 2016) which was attributed to discreet changes in HA antigenic sites (Benjamin S. Chambers et al., 2015; Stucker et al., 2015) involving a novel glycosylation site in the H3 hemagglutinin (HA) protein (Benjamin S. Chambers et al., 2015). Further sequence analysis, protein modeling and serology indicated that mutations in the HA and neuraminidase (NA) genes could be linked to escape from neutralizing antibodies (Broberg et al., 2015; Benjamin S. Chambers et al., 2015; Flannery et al., 2016; McCauley et al., 2015; Mostafa et al., 2016; Powell and Pekosz, 2020; Skowronski et al., 2016; Stucker et al., 2015; Tian et al., 2019).

It is critical for research and public health laboratories to have the methods in place for rapid reliable detection, recovery, and characterization of influenza seasonal drift variants. These capabilities inform decision making regarding vaccine composition and the public health response to an ongoing epidemic or pandemic (Flannery et al., 2016; Pebody et al., 2015).

In this report, we use influenza virus whole genome sequencing (WGS) and phylogenetic analysis to identify genetic variants, their frequencies, and their evolutionary relationships. We show that isolation of virus from nasopharyngeal swab (NPS) and nasopharyngeal wash (NPW) specimens using primary human nasal epithelial cell (hNEC) cultures is more efficient than isolation using a cell line. We found that amino acid changes in the NA (to introduce a glycosylation site near the sialidase pocket) and PB1-F2 (mutation H75P) altered virus replication and symptom severity during the 2014-15 influenza season. Furthermore, in this retrospective clinical cohort study, analysis of patient records showed that the co-existence of these NA and PB1-F2 genetic changes correlated with a higher incidence of cough and shortness of breath. Therefore, our data suggests that NA and PB1-F2 changes contributed to increased reporting of symptoms.

## Materials and Methods

### Sample collection and ethics

The human subjects protocol was reviewed and approved by the Johns Hopkins School of Medicine Institutional Review Board (IRB00052743). This protocol allowed for collection of residual nasopharyngeal swabs (NPS) from influenza positive patients from the Johns Hopkins Health System in the Baltimore-DC National Capital Region and collection of related, de-identified patient data from the electronic health records (EHR). The majority of specimens were from pediatric (under 18 years of age) and elderly (over 64) individuals, which represent the groups at increased risk for severe seasonal influenza disease (Luk et al., 2001; Morens et al., 2010; Skowronski et al., 2010). Symptoms were not reported for children under the age of two. Specimens from patients who had severe disease (as characterized by one of the following: supplemental oxygen use, Intensive Care Unit stay, or death). Specimens from age and sex matched individuals, who did not have characteristics of severe disease, were also included. Priority was also given to specimens from patients who had known influenza vaccination status documented in the EHR.

An IRB-exempt protocol allowed for access to residual nasopharyngeal wash specimens from de-identified patients of the Wright-Patterson Air Force Base (WPAFB). The WPAFB is the central repository for disease specimens from service men and women from around the country. A set of 16 samples that were collected before September from the WPAFB and grew to titers above 3 Log TCID50 in hNECs were selected for sequence analysis. Patient-level demographic and clinical data were not available for these specimens.

NPS and nasopharyngeal wash specimens were collected from each patient and placed in 3 ml of viral transport medium (MicroTest M4RT; Remel, Lenexa, KS, USA). All the specimens were aliquoted and stored at −70°C.

### Sample processing for genomic analysis

NPS samples were centrifuged for five minutes at 16,000xg to remove debris. Clarified supernatants were purified per manufacturer’s recommendations with the Ambion MagMax Viral RNA purification reagent, with the only protocol modification being omission of carrier RNA to the viral lysis enhancer reagent. RNA was resuspended in 20uL of supplied RNAse free elution buffer. Ambion Turbo DNA-free reagent (Life Technologies) was used to deplete genomic DNA. Manufacturer’s instructions for stringent DNAse treatments were followed: 2U of enzyme was added at the beginning of treatment and after 30 minutes of incubation at 37°C. Total duration of DNA depletion was one hour, after which DNAse inactivation reagent was added at 0.2 volumes.

Reverse transcription of viral RNA was carried out with Superscript III (Invitrogen) per manufacturer’s instructions for gene specific priming of low input samples using the Opti1 primer set, consisting of Opti1-F1 (GTTACGCGCC<u>AGCAAAAGCAGG</u>), Opti1-F2 (GTTACGCGCC<u>AGC</u>**<u>G</u>**<u>AAAGCAGG</u>), and Opti1-R1 (GTTACGCGCC<u>AGTAGAAACAAGG</u>). Samples were prepared using either a traditional multi-segment PCR approach (Zhou et al., 2009) or a preamplification-free approach, which we term amplification-limited viral RNA sequencing (vRNAseq). Briefly, primers and dNTPs were added to RNA and annealed by heating to 65°C for five minutes, then allowed to return to 4°C for five minutes. 400U of enzyme was then used in a total reaction volume of 40uL, and reverse transcription was carried out at 50°C. Upon completion, second strand synthesis was carried out at 16°C for 2.5 hours using mRNA Second Strand Synthesis reagent, which is comprised of *E. coli* DNA Pol I, E. coli DNA ligase, RNase H (NEB), in a total volume of 80µL. Double stranded viral cDNA was purified using one volume of Ampure XP per manufacturer’s instructions (Beckman Coulter). Once beads were completely dry, samples were eluted in 7.5µL of nuclease free water for five minutes prior to collection. One microliter was used for quantification with the High Sensitivity Qubit reagent according to manufacturer instructions (Invitrogen).

Nextera XT DNA library preparation was carried out per manufacturer’s instructions (Illumina). For samples that resulted in less than one nanogram or unquantifiable amounts of DNA, library preparation was carried out with the maximum amount of DNA available in 5uL. After tagmentation and addition of sequencing adapters by 12 cycles of PCR, samples were purified with one volume of Ampure XP and eluted in 10µL of nuclease free water. One microliter of the resulting library was then used for quantification with High Sensitivity Qubit reagents (Invitrogen), and multiplexed libraries were then pooled equi-mass, assuming an even size distribution between samples.

### Sequence data analysis

Raw reads from the Illumina MiSeq sequencer were first filtered using Trim Galore! [http://www.bioinformatics.babraham.ac.uk/projects/trim_galore/] using an adapter sequence of CTGTCTCTTATACACATCT and a quality score cutoff of 30. QC-trimmed reads were then individually classified using Kraken [https://github.com/DerrickWood/kraken] against a custom influenza-specific kraken database containing all complete sequences in EpiFlu with a modified taxonomy, which for flu types A, B, and C, identifies the likely segment, subtype, host, and year for each read. The most likely subtype for each sample was identified from the Kraken results, and 85 H3N2 samples were pulled out for further analysis. QC-trimmed reads for all 85 H3N2-identified samples were aligned to the A/Victoria/361/2011 reference sequence, and consensus FASTA sequences were generated for each sample. From these consensus FASTA sequences, 63 samples had at least 85% coverage of the HA segment and were used for subsequent phylogenetic analysis.

Reference sequences for the 2014-2015 influenza season were obtained from the GISAID EpiFlu database on October 22, 2016, by downloading all human H3N2 isolate sequences with a collection date between 2014-10-01 and 2015-05-31. In total, there were 5,642 unique EpiFlu ID numbers across 5,338 unique strain names. A subset of EpiFlu sequences was selected in two steps. First strains that contained exactly one of each of the 8 genome segments were selected, then for strain names that contained more than one EpiFlu ID number, only the strain with the lowest ID number was chosen. Sequences for the A/Victoria/361/2011 and A/Texas/50/2012 vaccine strains were obtained from NCBI BioProject PRJNA205638. The ultimate subset contained full genomes for 1,478 H3N2 strains. The same method was applied to select isolates for analysis from the 2013-2014 and 2015-2016 seasons, which led to 99 and 1140 isolates, respectively. The number of isolates from the 2013-14 season was an order of magnitude lower than the numbers for the other seasons because sequences for genes other than HA were often not available.

### Phylogenetic analysis

Data from each segment was combined into a segment-specific FASTA file for the 1,478 EpiFlu reference strains, the 63 sequenced samples with over 85% coverage in each segment, and the A/Texas/50/2012 vaccine strain, for a total of 1,542 isolates. Each segment was then aligned using MAFFT [http://mafft.cbrc.jp/alignment/software/], all deletions (‘-’) were replaced with ‘N’s, and all sequences were padded with ‘N’s to the length of the vaccine strain sequence. Any insertions with respect to the vaccine sequence were removed (any positions in the vaccine sequence that now contain ‘N’s), and all sequences were trimmed to the length of the vaccine sequence. Phylogenetic trees were created for each segment using the nucleotide option of FastTree [http://www.microbesonline.org/fasttree/], and trees were then re-rooted to the vaccine strain using the nw_reroot tool from the Newick utilities package [https://github.com/xflouris/newick-tools]. Trees were viewed and colored using FigTree [http://tree.bio.ed.ac.uk/software/figtree/] and HA clades were identified by adding known clade references to these trees and identifying branches for the 3C.2a, 3C.3, 3C.3a, and 3C.3b clades. From these branch identifications, a set of rules was developed (see below) so that a clade label could be added to each strain label. All clade information was added to the name of each strain name before tree generation, and then nodes were colored by searching for a specific clade text and coloring all nodes in that selection.

### Clade and genotype identification

Sequence data from each segment of the 1,542 uniquely named isolates with over 85% coverage in each segment were uploaded into a database using R. HA clades were identified using rules that selected sequences by amino acids (AAs) that associated with branches (https://hackmd.io/CwVgHAzAhgnAxgRgLQhiAZk4ATBMkwDsAbIUlFOsGMAEzHEAMEYQA===?view). The defining residues used were: 159Y for 3C.2a; 138S or 159S for 3C.3a; 157S for 3C.3b; 142R for 3C.2; 3L and 128A for 3C.3; and 3I and 128T for 3C.2. H3 numbering was used for all positions. Using these rules, all 1,542 strains were labeled according to the branch they fell on with the following exceptions: A/Brazil/9517/2015 is labeled 3C.3a but falls on the 3C.3b branch; A/Manitoba/RV3623/2014 and A/Nevada/30/2014 are labeled 3C.3b but fall on the 3C.3 branch; 01-13-P-0057 is labeled uncertain but falls on the 3C.2a branch; and 01-90-P-0102 is labeled 3C.3 but falls on the 3C.3b branch.

Consensus sequences of the twelve proteins PB2, PB1, PA, HA, NP, NA, M1, NS1 (primary open reading frames (ORFs)) and PB1-F2, PAX, M2, and NEP (alternative ORFs) were generated for each isolate that had over 85% sequence coverage and all of the reference strains. Sequences of the eight primary ORF proteins were generated by translation from the first ATG in each gene to the first stop codon. Sequences of the four alternative ORF proteins were generated with the following set of rules: translation of PB1-F2 was programmed to start at the 4^th^ ATG of segment 2 and stop at the first in frame stop codon; translation of PAX started from the first start codon in segment 3, included the first 573 nucleotides, then shifted +1 to nt 575 and carried on to the next stop codon in that frame; M2 was translated from the first start codon of the segment 7 through 51 nts, joined to nt 715 and stopped at the first stop codon in frame 2; NEP was translated from the first start codon in segment 8 through the 30^th^ nt, joined with base 503 and stopped at the in-frame stop codon.

Genetic variation among clade 3C.2a isolates was analyzed further. Amino acid alignments were generated using MAFFT and the frequency of each AA residue at every location was determined using Excel (Microsoft). Instances where the consensus AA fell below 75% frequency were noted. Mutations associated with putative NA glycosylation signatures were identified by the residues at positions 245-247. Isolates were labeled as NAg+ if those residues were NAT or NAS and NAg-for all other sequences. Residues at AA position 75 within PB1-F2 were also identified for each isolate.

### Cell culture and serum free media

Madin-Darby canine kidney cells (MDCK) were cultured as previously described (Grantham et al., 2010). MDCK-SIAT cells were kindly donated by Scott Hensley and cultured exactly the same as MDCK cells. Cultures of human primary, differentiated nasal epithelial cells (hNEC) were obtained from volunteers and differentiated at an air-liquid interface (ALI), as previously described (Fischer et al., 2015; Forero et al., 2017; Ramanathan et al., 2009; Wohlgemuth et al., 2017).

### Virus isolation

Virus isolation with MDCK cells was performed by incubating 100µl of NPS on twice-PBS-washed cells in 48-well plates for one hour at 32°C with 5% CO_2_ with agitation every 15 minutes. After inoculation, NPS media was aspirated and cells were incubated again at 32°C with 5% CO_2_ in infection media (Dulbecco’s Modified minimal media; 0.3% bovine serum albumin) with 5 µg/ml N-acetyl trypsin (NAT). Media was collected and replaced daily.

Virus isolation with hNEC cultures was performed by incubating 100µl of NPS on the twice-PBS-washed apical surface of hNEC cells in 12-well transwell plates (Corning) for two hours at 32°C with 5% CO_2_. Inoculum was removed and cells were returned to incubate at 32°C with 5% CO_2_ for up to 7 days. The hNEC cultures were maintained with media on the basolateral side, which was changed every 2-3 days. Apical washes, acquired by adding 300ul infection media to the apical surface of hNECs and incubating for 10 minutes at 32°C with 5% CO_2_, were collected daily. All samples were stored at −70°C.

### Infectious virus quantification

The infectious virus titers were determined by calculating the 50% tissue culture infectious dose (TCID_50_) using the Reed and Muench method (McCown and Pekosz, 2005; Reed and Muench, 1938).

### Virus generation and growth analysis

Infected-hNEC culture supernatants, from 4 dpi of hNECs, were inoculated into MDCK cells with a multiplicity of infection (MOI) of 0.01 or 0.001 TCID_50_ units per cell and incubated in infection media with 5µg/mL NAT for 5 days at 32°C to make virus stocks. A/Columbia/P0041/2014 (Co14) has the NAg-F2H genotype while A/Bethesda/P0055/2015 (Be15) has the NAg+F2P genotype. Infectious virus was quantified by TCID_50_. HA, NA, and PB1 genes of amplified virus stocks were Sanger sequenced. The sequences of virus stocks were compared to the WGS consensus sequences and the HA of both Co14 and Be15 were found to have the non-synonymous nucleotide mutation C478A, which encoded the mutation T160K (H3 Numbering). This mutation resulted in the loss of a predicted glycosylation site at position 158 in the HA protein.

For MDCK and MDCK-SIAT cell growth curves, cells in 24-well plates were washed twice with PBS containing Ca^2+^ (0.9mM) and Mg^2+^ (0.5mM) (Sigma) and infected with MOI 0.001 virus, diluted in 100 µL of infection media. After 1 hour inoculation, the supernatant was removed, cells washed 3 times with PBS with calcium and magnesium and incubated in infection media with 2.5 µg/mL NAT at 32°C with 5% CO_2_. Samples were collected at indicated times by removing and replacing the media. Supernatants were stored at −70°C and titrated by TCID_50_.

For hNEC growth curves, the apical surface of 24-well hNEC cultures was washed twice with PBS and infected with MOI 0.01 virus in 100 µL in infection media. hNECs were incubated with the inoculum for 2 hours at 32°C with 5% CO_2_. Inoculum was removed and the apical surface was washed twice with PBS before cells were returned to the incubator. Apical wash samples were collected, at indicated times, by incubating 100 µL of infection media on the apical surface for 10 minutes at 32°C as previously described (Fischer et al., 2015). Supernatants were stored at −70°C and infectious virus quantified by TCID_50_.

Each virus was inoculated into 4 wells of each cell type and growth curves were performed on two separate occasions, for a total of 8 wells per virus, per cell type, per time point. The average viral titers from the 8 equally treated wells were graphed for virus comparisons.

### Plaque assay

MDCK cells, at approximately 100% confluence were washed twice with PBS with calcium and magnesium. Virus was serially diluted 10-fold in infection media and 250 µL of each dilution was added to cells. The plates were incubated for one hour at 32°C in 5% CO_2_ with agitation, then inoculum was removed and cells were overlaid with 1X MEM with the addition of 0.3% BSA, 10mM Pen/Strep, 5mM Glutamine, 1% agarose (Invitrogen), 1 mM Hepes (Gibco), and 5 µg/mL NAT and incubated at 32°C in 5%CO_2_ for 72 hours, after which cells were fixed with 4% paraformaldehyde (Fisher) and stained with Napthol Blue Black (Sigma). One image per well was collected using an Olympus OM-D E-M5 Mark II digital camera. The scale for all images was determined by photographing a ruler. Images were processed in ImageJ v1.49 Plaque images were cropped to remove area outside of the well, then they were turned into 8-bit images. Next, plaques were circled manually using the selection tool. Finally, plaques were analyzed using the ImageJ analyze particle size command with a size minimum of 200 pixels and a circularity of 0.10-1.00. In total 40-60 plaques (between 10-20 plaques per well) were analyzed per virus.

### Plasmids

Influenza A/Bethesda/55/2015 (Be15, ID 253812) virus was isolated on hNECs and expanded for one passage on MDCK cells to make a working stock. Viral RNA was extracted from the working stock using the Qiagen viral RNA purification kit. PB1-F2 (75P) cDNA was amplified via one step RT-PCR using SuperScript III/Platinum Taq mix (Invitrogen). The RT-PCR product was cloned into pCAGGS mammalian expression vector via restriction enzymes KpnI and SphI. A C-terminal FLAG epitope tag (DYKDDDDK) was added via site-directed mutagenesis (QuikChange Lightning, Agilent). The 75^th^ amino acid in the PB1-F2 protein was changed to a histidine (H) via site directed mutagenesis (QuikChange Lightning, Agilent) to produce the PB1-F2-75H plasmid. The NA-FLAG pCAGGS plasmids for A/Bethesda/55/2015 (ID 253812, NAg+) and A/Columbia/41/2014 (Co14, ID 253817, NAg-) were generated as previously described (Powell and Pekosz, 2020).

### Cell transfection and lysate preparation

For PB1-F2 localization studies, A549 cells were cultured in complete medium. Cells were grown in 12 well chamber slides (Ibidi). For PB1-F2 expression, cells were transfected with 0.5µg of DNA and 4µl of Lipofectamine 2000 reagent (Invitrogen) per well. Transfection was allowed to proceed for four hours then media was replaced with fresh complete medium. Twenty-four hours post transfection, complete media was removed, the cells were washed once in PBS, then fixed using 4% paraformaldehyde (Sigma). Cells were stored in PBS at 4ºC until processed for microscopy. For expression of NA proteins, HEK293T cells were grown in a 6-well plate in complete medium. 2.5µg of pCAGGS NA-FLAG (either NAg+ or NAg-) were transfected using 7.5µl of LT1-TransIT transfection reagent (Mirus). Twenty-four hours post transfection, complete medium was removed and cells were lysed with RIPA buffer for immunoprecipitation of FLAG tagged NA protein, as described below. Cell lysates were stored at −20ºC.

### Microscopy and image analysis

A549 cells transfected with PB1-F2 constructs (described above) were fixed in 2% paraformaldehyde for 15 minutes at room temperature and then permeabilized with 0.2% Triton X-100 (Sigma) in PBS for 15 minutes before blocking. Blocking buffer consisted of PBS with 2% normal donkey serum (Sigma) and 0.5% BSA (Sigma). Cells there then incubated with polyclonal rabbit anti-Tom20 (sc-11415; Santa Cruz) (2µg/ml) and monoclonal anti-FLAG (clone M2; Sigma) (10µg/ml) antibodies. After primary antibody incubation, cells were washed with 0.2% Tween 20 in PBS. Next, cells were incubated with secondary antibodies Alexa Fluor 488 donkey anti-mouse IgG (Invitrogen) (4µg/ml) and Alexa Fluor donkey anti-rabbit IgG (Invitrogen) (4µg/ml). Cells were mounted in Prolong Gold anti fade with DAPI and cover slipped. Cells were imaged using a Zeiss LSM 700 microscope with Zen software for image acquisition. Images were taken with a 100x/1.4 Oil DIC objective in Z-Stack mode with 0.5µM space between stacks. Images were analyzed and process with Volocity and FIJI software. Individual cells were manually identified and then M2 colocalization coefficients were calculated between PB1-F2 FLAG (red) and Tom20 (green) using Volocity. Statistics were performed in Prism (Graphpad) with a Welch’s t-Test, *p<0.05.

### Immunoprecipitation and PNGaseF treatment

FLAG-tagged neuraminidase proteins were immunoprecipitated from cell lysates per the manufacturer’s instruction (Abcam). Briefly, cell lysates were made using RIPA buffer and allowed to lyse at 4ºC for 1 hour. After the extended lysis step, lysates were clarified by centrifugation at 10,000g for 10 minutes to precipitate cellular debris. The supernatant from this clarification step were incubated with mouse anti-FLAG antibody (Sigma, clone M2 1µg/ml) over night at 4ºC with gentle rocking. The following day, washed protein A/G coupled agarose beads were added to the cleared lysate and incubated with rocking for 2 hours at room temperature. Afterwards, the beads and lysate mix were centrifuged at 3,000g for 3 minutes to pellet immunoprecipitation beads. The immunoprecipitation beads were washed three times with kit wash buffer. Next, 100µl of 1% SDS in PBS was added and mixture was boiled at 100ºC for 10 minutes. The mixture was centrifuged again at 3,000g for 3 minutes and the supernatant containing precipitated NA protein was used for PNGase treatment. PNGase F treatment was done according to the manufacturer’s instructions (NEB).

### Western Blotting

After PNGaseF treatment, samples were mixed with 4X-Laemli buffer (Bio-Rad) containing 250 mM dithiothreitol (DTT, ThermoFisher Scientific) and boiled at 100ºC for 5 minutes. Samples were run on 4-20% Mini-PROTEAN TGX gels (Bio-Rad) with an All-Blue precision plus protein ladder (Bio-Rad) at 70 V for approximately 1 hour. Proteins were transferred onto an immobilon-FL membrane (Millipore) at 75 V for 1hr. After transfer, membranes were blocked with blocking buffer for 1 hour at room temp on a rocker. Blocking buffer consisted of PBS containing 0.05% Tween-20 (Sigma) and 5% non-fat milk (Bio-Rad) for 1 hour at room temp. Rabbit anti-FLAG antibody (Cell Signal Technology, #2368, 0.5µg/ml, diluted in blocking buffer) was added and allowed to bind overnight at 4ºC in blocking buffer. Membranes were washed in PBS with 0.05% Tween-20 (wash buffer). Secondary antibody was added for 1hr at room temperature (25ºC) in blocking buffer then washed again in wash buffer. Blots were imaged and analyzed with the FluorChem Q system (Proteinsimple).

### Statistical analysis

Differences between the average viral titers of isolates grown on MDCK cells or hNECs were determined by paired, two-tailed, t-test. Virus replication differences in low MOI growth curves were determined with repeated measures two-way ANOVA followed by a Bonferonni post-test. A multivariate logistic regression analysis was performed in R to determine whether or not any co-morbidities associated with genotype differences and disease symptoms.

## Results

### Sequence characterization of virus

Ninety-four influenza positive NPS specimens with demographic and clinical data were collected from three hospitals in the Johns Hopkins Health System, referred to here as Johns Hopkins Medical Institutes (JHMI). Sixteen influenza positive NPW specimens that lacked demographic and clinical data were provided by the Wright Patterson Airforce Base (WPAFB). Whole genome sequencing (WGS) was performed on 77 of the JHMI and all 16 of the WPAFB specimens using an amplification strategy designed to specifically detect influenza genomic sequences. Illumina sequencing and Kraken analyses were used to generate and categorize consensus sequences for the eight influenza virus gene segments in each specimen.

HA coding region sequence reads were used to determine the influenza virus type 86% (66/77) of the JHMI samples and 88% (14/16) of the WPAFB samples (Table 1). The influenza subtype was determined for 84% (65/77), and the HA clade was determined for 77% (59/77), of JHMI isolates. The influenza subtype and clade were determined for 88% (14/16) of the WPAFB samples.

**Table 1.** Influenza virus classification by WGS. A total of 77 JHMI and 16 WPAFB nasal swab samples were sequenced and influenza virus type, subtype, and clade were determined.

| Influenza |  | WGS JHMI | WGS WPAFB |
| --- | --- | --- | --- |
| Type A |  | 86% (66/77) | 88% (14/16) |
| Subtype H3N2 |  | 84% (65/77) | 88% (14/16) |
| Clade | (determined) | 77% (59/77) | 88% (14/16) |
|  | 3C.2 | 0% (0/59) | 0% (0/14) |
|  | 3C.2a | 93% (55/59) | 57% (8/14) |
|  | 3C.2b | 0% (0/59) | 0% (0/14) |
|  | 3C.3 | 0% (0/59) | 7% (1/14) |
|  | 3C.3a | 5% (3/59) | 14% (2/14) |
|  | 3C.3b | 2% (1/59) | 21% (3/14) |

### HA clade distribution

The distribution of HA clades differed between WPAFB and JHMI samples, with 57% of WPAFB and 94% of JHMI samples belonging to H3N2 clade 3C.2a (Figure 1 A, B). The remaining isolates were characterized as either 3C.3a or 3C.3b, with one WPAFB isolate characterized as a 3C.3. Clade diversity among the WPAFB viruses was higher compared to the JHMI isolates (Figure 1 A, B). WPAFB samples were collected earlier in the 2014-15 influenza season therefore the timing of collection likely accounts for the differences in clade distribution between these two populations.

**Figure 1.**
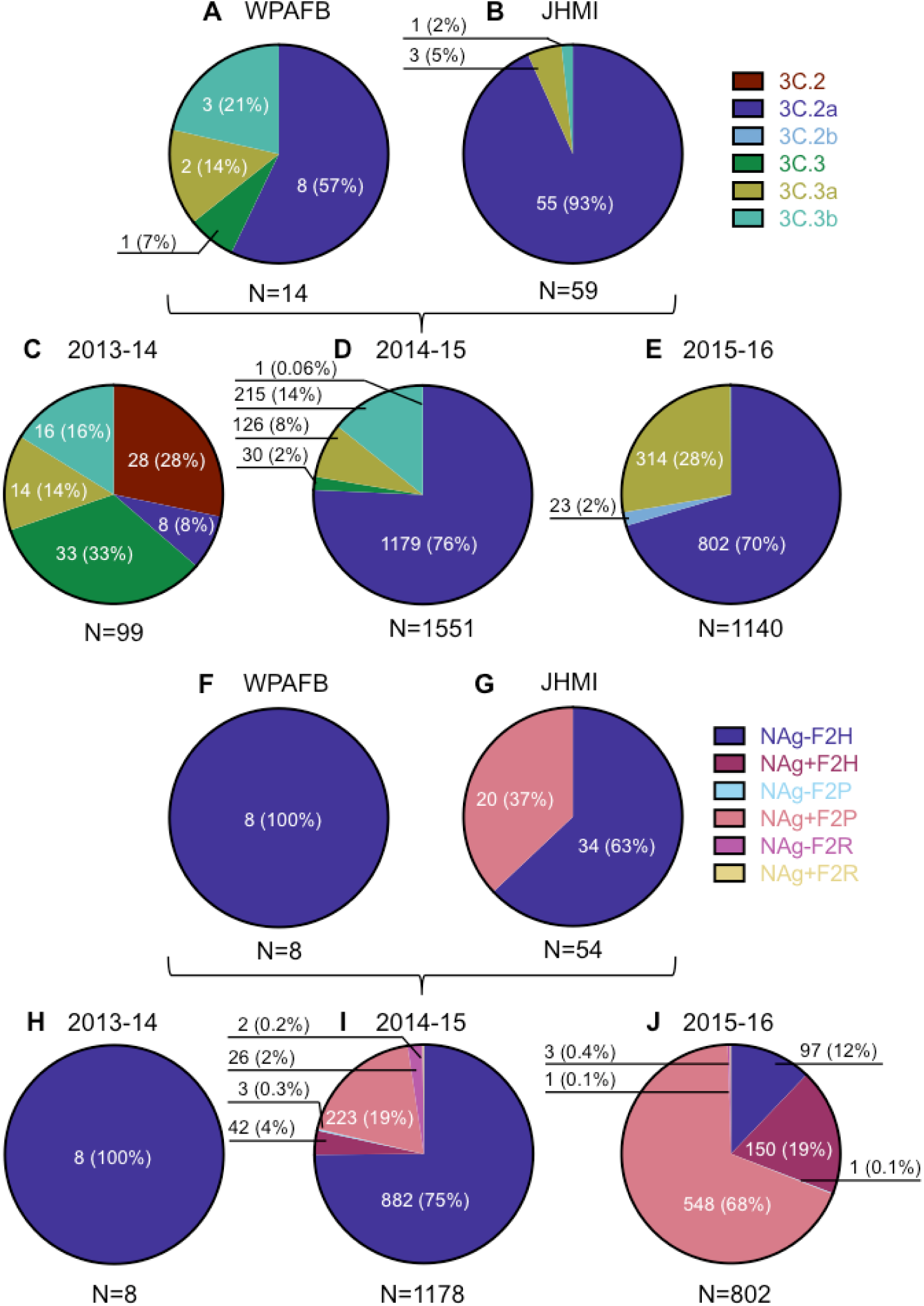
Characterization of human isolates by clade and genotype. The number of virus isolates that belong to clades and genotypes are reported according to sequence only. The prevalence of each clade is represented in pie charts A-E. The prevalence of each 3C.2a genotype, as a percentage of clade 3C.2a isolates, is represented in charts F-J. Pie charts A and F represent isolates from WPAFB, charts B and G represent isolates from JHMI. EpiFlu isolates from the 2013-14 season are represented in C and H and those from the 2015-16 season are represented in E and J. The pie charts for 2014-15 include both EpiFlu sequences and the sequences from this study.

For context, the HA clade distributions of viral isolates with full genomes published on the EpiFlu database were determined for the northern hemisphere winter seasons of 2013-14, 2014-15 and 2015-16. H1N1 dominated during the 2013-14 season, so H3N2 isolates were less frequent and interestingly, their clade distribution was diverse (Figure 1 C). H3N2 viruses dominated during the 2014-15 influenza season and clade 3C.2a viruses were by far most common, as was previously reported (Flannery et al., 2016). JHMI and WPAFB sequences were included in the analysis for the 2014-15 season for a total of 1551 isolates, of which 76% (1179) belonged to clade 3C.2a (Figure 1 D). H3N2 and 3C.2a clade viruses also dominated during the 2015-16 season (Figure 1 E).

### Identification and frequency of NA glycosylation and PB1-F2 amino acid 75 mutations

Further analysis of the influenza virus sequences revealed that over one third (37%; 20/54, Figure 1 G) of JHMI isolates had a non-synonymous nucleotide mutation (ntG734A; open reading frame (ORF) numbering throughout) in the NA segment. This mutation changed a Serine (S) residue to Asparagine (N), thereby creating a putative N-linked glycosylation signal in NA at amino acid residues 245-247 (NAg+). In addition, 18 of the 20 isolates had another non-synonymous mutation (ntT739A), which changed the Serine residue to Threonine (T) at amino acid 247. Since the putative glycosylation sequence is N-X-S/T, the second non-synonymous mutation in NA presumably would not affect the glycosylation state at this site. This glycosylation site was previously observed in about a third (27%; 3/11) of 2014-15 influenza virus isolates (Mostafa et al., 2016). A glycosylation motif at this site reduced enzyme activity, reduced or abolished antibody binding, and reduced antibody-based protection against influenza virus infection in mice (Powell and Pekosz, 2020; Wan et al., 2019b). Interestingly, the WPAFB samples did not contain the NAg+ mutations (Figure 1F). Since these samples were collected earlier in the 2014-15 season, the data may indicate that the NAg+ mutation emerged later in the season.

The only other non-synonymous mutation that occurred with greater than 25% frequency among the 3C.2a clade JHMI isolates was an ntA318C mutation in the PB1 ORF, which resulted in a change from Histidine (H) to Proline (P) at amino acid 75 (H75P) of the PB1-F2 ORF (F2P). The same 20 JHMI 3C.2a clade viruses that were NAg+ contained the F2P mutation while all other 3C.2a isolates lacked both mutations (NAg-F2H) (Figure 1 G). None of the 3C.2a clade isolates from the WPAFB had the F2P genotype (Figure 1 F). This data indicated that the NAg+ and F2P genotypes were co-selected in clade 3C.2a viruses.

All of the clade 3C.2a influenza winter season isolates from 2013-2016 that had WGS were characterized into six genotypes. These were determined by the presence or absence of a glycosylation motif at residues 245-247 in the NA protein and the amino acid residue at position 75 in the PB1-F2 protein (Figure 1H-J). Besides Histidine and Proline, the only other amino acid that was present at position 75 in PB1-F2 was Arginine (R). In samples from 2013-14, only the NAg-F2H genotype was detected (Figure 1H). The NAg-F2H genotype was dominant during the 2014-15 winter season (75%) with the NAg+F2P genotype being the second most common genotype (19%) (Figure 1I). The frequency of NAg+F2P isolates grew to 68% of all clade 3C.2a isolates in 2015-16 while the NAg-F2H genotype shrank from 75% to 12%. The only other genotype to increase in frequency was NAg+F2H, which grew from 4% to 19% (Figure 1I and J). Taken together, the data indicate that the NAg+F2P genotype rapidly became dominant among clade 3C.2a viruses (compare Figure 1I to J), while the frequency of 3C.2a viruses remained relatively similar (compare Figure 1D to E) between the 2014-15 and 2015-16 seasons. Viruses with NAg+ accounted for 87% of isolates in the 2015-16 season showing that NAg+ viruses were more frequent than viruses with the PB1-F2P mutation (Figure 1J).

### Evolution of NA glycosylation and PB1-F2_75P

HA, NA, and PB1 phylogenetic trees with the 1487 2014-15 winter season EpiFlu, JHMI, and WPAFB isolate sequences were generated to determine the phylogenetic relationships between genotypes (Figure 2). HA clades 3C.2a, 3C.3, 3C.3a, and 3C.3b were all represented in the HA phylogenetic tree (Figure 2A). The branches of the NA and PB1 phylogenetic trees use the HA clade color scheme, with the exception of viruses encoding the NAg+ or F2P mutations, which use the 3C.2a genotype color scheme from Figure 1 (salmon and burgundy in the NA and PB1 trees, respectively, figure 2B,C). In the NA phylogenetic tree, the large set of salmon colored branches at the top indicates that most detected NAg+ isolates arose from a common ancestor. In the PB1 phylogenetic tree, the large set of burgundy colored branches at the top similarly indicates that all detected F2P isolates arose from a common ancestor.

**Figure 2.**
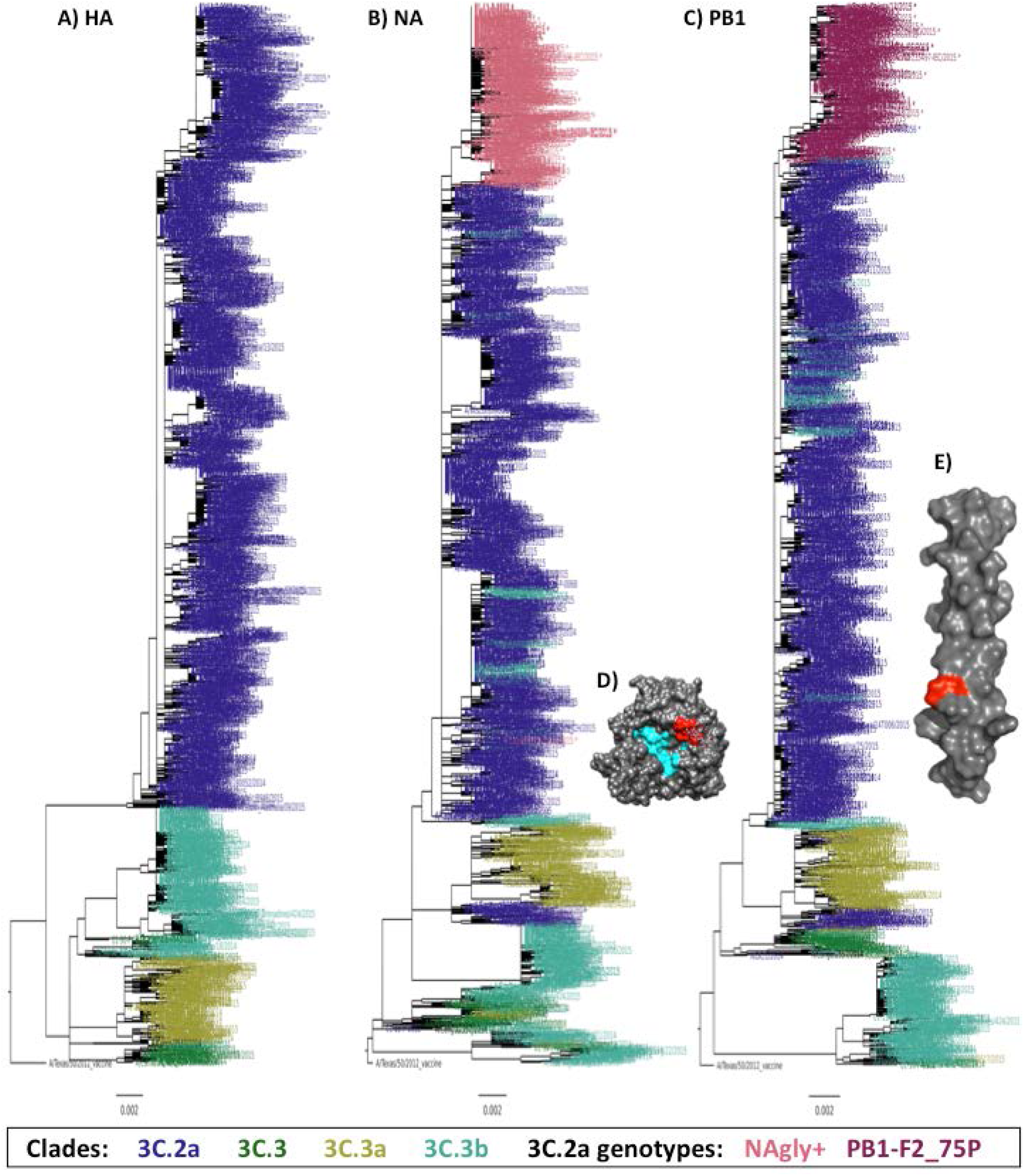
HA, NA, and PB1 phylogenetic trees and crystal structures. (A, B, C) Each tree was made with sequences from the gene segment indicated. The sequences included 1116 from the EpiFlu database that had full length nucleotide coverage for all open reading frames isolated from 10/01/2014 to 5/31/2015 plus 54 JHMI and 8 WPAFB isolates for a total of 1178 genomes. Phylogenetic trees were made using FastTree and images were made with FigTree. Branches are colored by clade (HA) or clade and genotype (NA and PB1) as indicated in the key. (D) X-ray diffraction crystal structure of NA from A/Tanzania/205/2010 H3N2 (PDB 4GZP). The enzymatic site is colored cyan and the amino acids that encode the putative glycosylation site of NAg+ viruses are colored red. (E) Solution NMR structure of PB1-F2 C-terminal domain AA residues 50-87 from A/Puerto Rico/8/1934 (PDB 2HN8). Residue 75 is colored red. (D,E) Visualized using UCSF Chimera v1.11.1.

Mutations were mapped onto protein structures to determine their location. The putative NA glycosylation site was mapped to the X-ray crystal structure of the NA protein from influenza A/Tanzania/205/2010 (H3N2) (Zhu et al., 2012). It showed that the site was on the surface of the NA protein and directly adjacent to the NA enzymatic site (Figure 2D). The location of PB1-F2 residue 75 was mapped to the NMR structure of the C-terminus tail of the PB1-F2 protein from influenza A/Puerto Rico/8/1934 (H1N1)(Bruns et al., 2007). It shows that the mutation site is about two-thirds of the way down the C-terminal domain (Figure 2E).

Since the NAg+ and F2H genotypes appear together in most virus strains (Figure 1I and J), it is likely that one common ancestor led to the dominance of this genotype in 2015-16 clade 3C.2a viruses. To determine the genetic relationships between the JHMI and WPAFB viruses, we performed a phylogenetic analysis on concatenated genome segments from the JHMI and WPAFB samples (Figure 3A). The NAg+F2P genotype viruses all clustered together from a single branch of the tree (salmon color at top of tree), indicating that a common ancestor acquired both the NAg+ and the F2P mutations. To determine if this result was the same in the general population of viruses we performed a similar phylogenetic analysis on all H3N2 viruses with full genome sequences available from the 2014-15 and 2015-16 seasons (Figure 3B). Again, the NAg+F2P viruses clustered together providing further support for the presence of a single common ancestor from which all the NAg+F2P genotype viruses derived.

**Figure 3.**
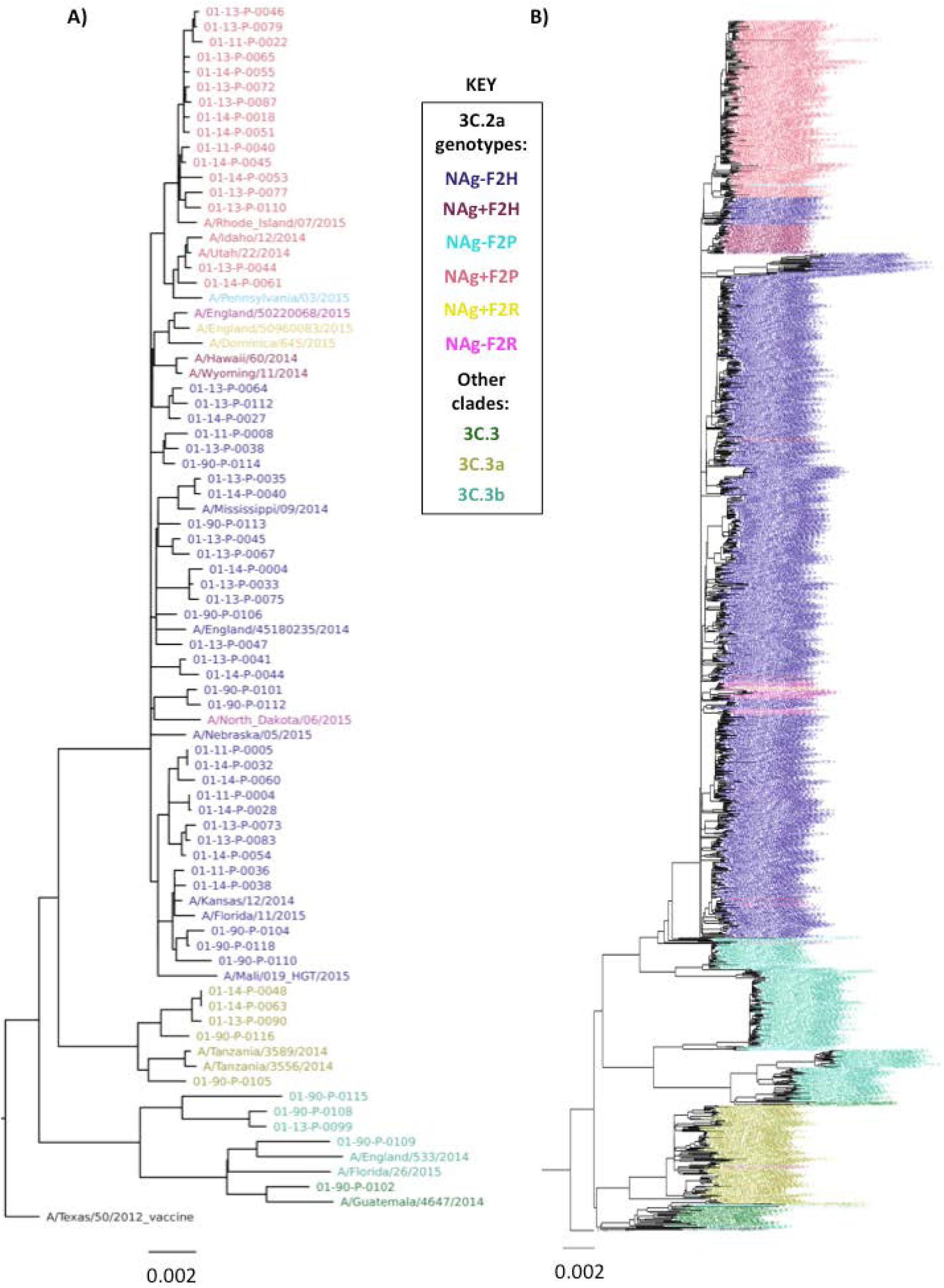
Phylogenetic trees of concatemerized influenza virus genomes. (A) Sequences from the 54 JHMI, and 8 WPAFB isolates and some reference strains were concatenated and used to construct a phylogenetic tree. (B) A second phylogenetic tree of concatenated genes was generated using the 1178 2014-15 isolates used in Figure 2 plus all 2015-16 isolates from the EpiFlu database. Trees were made with the same methods for Figure 2. Branches are colored by clade or genotype as indicated in the key.

The locations of the NAg-F2P viruses (cyan branches) within the NAg+F2P viruses (salmon branches) indicate that the NAg-F2P genotype likely arose from mutations that reversed the NAg+ to the NAg-genotype in the NAg+F2P genotype lineage. There was no evidence that the F2P mutation was gained by NAg-viruses. The group of NAg+F2H viruses (plum) that branched from the trunk of the tree before the NAg+F2P group, and the other NAg+F2H virus branch locations, scattered among NAg-F2H branches, suggest that the NAg+ mutation was selected numerous times and that in one case, it was selected slightly before selection of the H75P mutation. This indicates that the H75P mutation was only selected after NAg+ was present.

NAg+F2R isolates seem to have been selected in the opposite order. The branching of the three NAg+F2R viruses (yellow) among a small group of NAg-F2R viruses (magenta) indicates that a subset of F2R viruses created a background in which NAg+ was selected, but this group was short-lived and did not lead to the emergence of the NAg+F2P genotype. In the PB1 phylogenetic tree all F2P and all F2R viruses shared a single node (Figure 2C), indicating that, evolutionarily, these genes were closely related.

The two most important points from the phylogenetic analysis are that (1) the two mutations of the NAg+F2P genotype clade 3C.2a viruses descended from a common ancestor and (2) expansion of NAg+F2P genotype viruses suggests selection according to a fitness advantage of that genotype over other viruses within the 3C.2a clade.

### Virus isolation on hNEC cultures was more successful than on MDCK cells

Virus isolation was attempted from 104 of the JHMI and WPAFB NPS specimens. Primary differentiated hNEC cultures were inoculated with all 104 NPS specimens while MDCK cells were inoculated with a subset of 96 specimens. Specimen use was dictated by available sample volume. Infectious virus titers were determined by TCID_50_ assay for supernatants collected from hNEC or MDCK cells at four days post infection (dpi). Inoculation of hNEC cultures led to isolation of viruses from 69 (66%) of the NPS specimens while inoculation of MDCK cells only led to isolation of viruses from 22 (23%) of the specimens (Figure 4A). Isolation using hNEC cultures had a nearly three-fold higher success rate. The median infectious virus titers were 3.6 log_10_ TCID_50_/ml from hNEC cultures and 2.4 log_10_ TCID_50_/ml, indicating the clinical isolates grew to higher infectious virus titers in hNEC cultures compared to MDCK cells (Figure 4A). The higher virus isolation frequency and higher viral titers achieved with hNEC cultures were consistently observed, irrespective of the HA clade or genotype of virus isolated (Figure 4B). This data indicates that primary hNEC cultures, which more accurately represent the cells that influenza viruses encounter in the upper respiratory tract, are more efficient than MDCK cells at isolating virus from clinical specimens.

**Figure 4.**
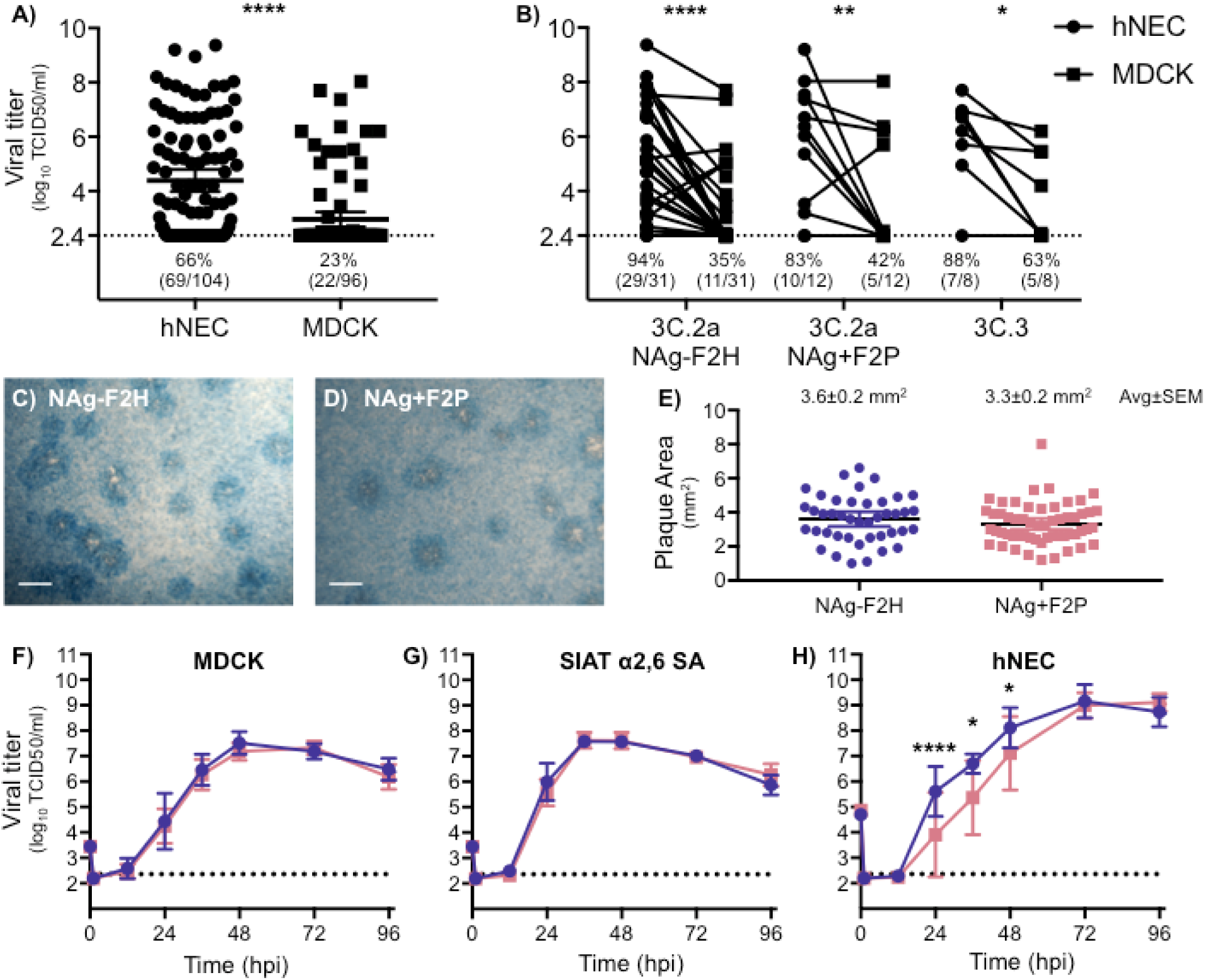
H3N2 virus isolation and characterization. (A) Virus isolation was attempted with 104 NPS specimens on hNEC cultures and 96 NPS specimen on MDCK cell cultures. Apical washes and supernatants were collected from hNEC and MDCK cell cultures, respectively, at 4 dpi and infectious virus titers determined by TCID_50_. Lines represent means and error bars represent 95% CIs. Statistical significance was determined with a Wilcoxon matched pairs signed rank test. (B) Comparison of hNEC and MDCK 4dpi viral isolation titers was performed for three major genotypes. The percent and ratio of viruses that grew to levels above the limit of detection are noted below each dataset (A,B). Repeated Measures two-way ANOVA indicated that virus had no significant effect on growth and growth in MDCKs was significantly lower than that achieved in hNECs for all three virus genotypes. The limit of detection was 2.36 log_10_ TCID_50_/ml. (C-E) Plaque assays were performed by inoculating MDCK cells and incubating for 3 days, under agarose, at 37ºC. Representative images of plaques are shown for each virus (C,D). The area of over 40 plaques for each virus was determined (E) and analyzed using ImageJ. The average of plaque areas between the two viruses was determined by a two-tailed unpaired t-test. (F-H) Viruses isolated on hNECs and expanded once in MDCK cells were used to compare viral growth in MDCK cell (F), MDCK-SIAT cell (G), and hNEC (H) cultures. Cultures were inoculated with MOI 0.001, 0.001, and 0.01, respectively. Supernatants or apical washes were collected at times indicated and infectious virus determined by TCID_50_. N=8 wells per virus, per cell type, per time point. Error bars represent SD. Statistical significance was determined with a Repeated Measures two-way ANOVA with Bonferroni posttest correction. *P<0.05, **P<0.01, ***P< 0.001, ****P<0.0001

### Virus growth and plaque morphology

Viruses representative of the two 3C.2a genotypes – A/Columbia/P0041/2014 (NAg-F2H) and A/Bethesda/P0055/2015 (NAg+F2P) – were expanded and assessed for differences in replication on three different cell types: MDCK cells, MDCK-SIAT cells, and hNECs. MDCK-SIAT cells overexpress the α2,6-linked sialic acid utilized by human influenza viruses as a cellular receptor (Matrosovich et al., 2003). Plaque assays were performed on MDCK cells and the two genotypes showed similar plaque morphologies (Figure 4C and D) and areas (Figure 4E). In low multiplicity of infection (MOI) growth curves, the two viruses grew to similar titers on MDCK cells (Figure 4F) and MDCK-SIAT cells (Figure 4G).. The NAg-F2H virus grew to higher titers on hNEC cultures compared to NAg+ F2P (Figure 4H) and this difference was statistically significant at 24, 36, and 48 hpi. The replication difference in hNECs was transient and both viruses reached equivalent peak titer at the same day post infection (72 hpi). Taken together, the data indicate that the two genotypes replicate to similar extents in immortalized cell lines but show different replication kinetics in hNEC cultures.

### Characterization of PB1-F2 and NA proteins

Plasmids encoding the PB1-F2-75H and PB1-F2-75P genes were transfected into immortalized A549 cells to study differences in intracellular localization. Twenty-four hours after transfection, cells were fixed and imaged for PB1-F2 and Tom20, a protein that localizes to the outer membrane of the mitochondria (Yamamoto et al., 2011) (Figure 5A). The PB1-F2-75H was found to have significantly higher colocalization with the mitochondria protein compared to PB1-F2-75P (Figure 5B). This suggests that the 75^th^ amino acid in PB1-F2 impacts the ability of this protein to localize to mitochondria.

**Figure 5.**
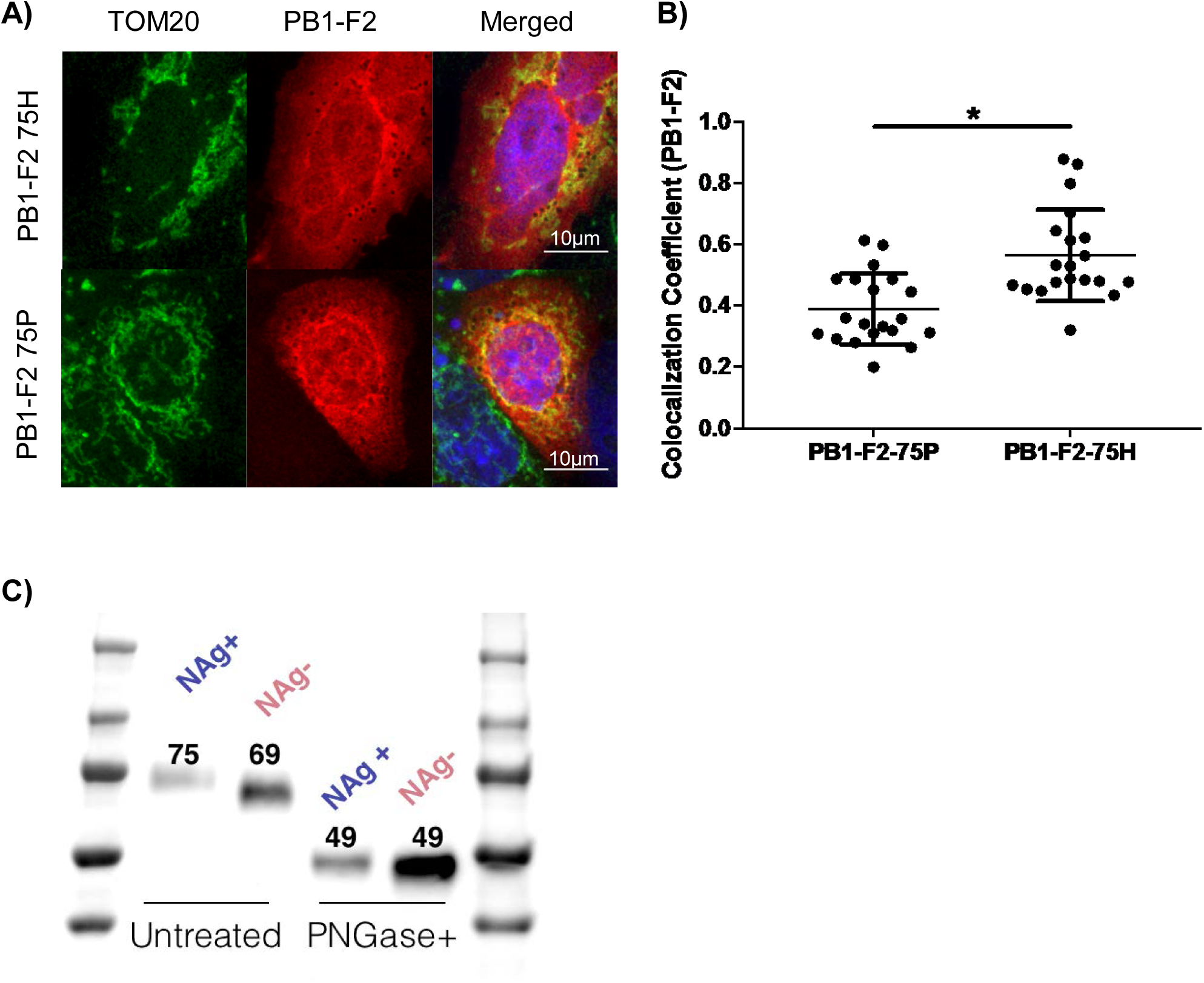
PB1-F2 localization and Western blot analysis of NAg+/- proteins. (A) Representative microscopy image of A549 cells that were transfected with a plasmid containing a FLAG epitope tagged PB1-F2-75P or PB1-F2-75H. Twenty-four hours post transfection, cells were fixed and stained for the mitochondrial marker Tom20 (red) and the FLAG epitope tag (red) or nuclei (DAPI). (B) Analysis of colocalization of PB1-F2 protein and mitochondrial marker Tom20. Coefficient of colocalization with Tom20 was analyzed in Volocity from 20 cells transfected with PB1-F2 75P or PB1-F2 75H. Statistical significance was determined in Graphpad Prism with a Welch’s t-Test, *p<0.05. (C) HEK293T cells were transfected with plasmids containing the FLAG epitope tagged NAg+ or NAg-gene, immunoprecipitated, treated with PNGaseF then analyzed via SDS-PAGE and western blot. Numbers above protein bands indicate approximate molecular weight based on protein ladder (left and right of samples).

To verify that the putative N-linked glycosylation site on NA is being utilized, plasmids encoding the cDNAs for either NAg- and NAg+ were transfected into immortalized HEK293T cells and analyzed for molecular weight via SDS-PAGE and western blot. The NAg+ protein had a molecular weight of 75kDa and the NAg-protein had a molecular weight of 69kDa. This size difference is indicative of an additional N-linked glycan on the NAg+ protein. When both NA proteins were treated with PNGaseF to remove all N-linked glycans, the two proteins migrated at 49kDa, which is the predicted molecular weight of the human H3N2 NA protein. These results suggest that this putative glycosylation site is accessible by host cell glycosylation machinery and is glycosylated. This glycosylation site, at position 245, alters the antigenicity of the protein as well as significantly decreases viral replication in primary respiratory epithelial cells (Powell and Pekosz, 2020; Wan et al., 2019a).

### Clinical characteristics of patients

Clinical and demographic characteristics were retrospectively analyzed for the 94 JHMI influenza positive NPS specimens (Table 2). There were 54% females and 46% males. The racial group distribution was 46% white, 29% black and 11% Asian. Vaccination status was known for 36% of patients, of which 73% were vaccinated. 61% of patients were treated as outpatients and 10% required inpatient care. 30% of patients required intensive care and most of those (97%) required supplemental oxygen. 5% of the patients died. 57% of patients had one or more co-morbidities. The JHMI samples were supplemented with 16 nasal wash specimens from the Wright-Patterson Air Force Base (WPAFB) repository. Clinical data for the WPAFB patients was not accessible.

**TABLE 2:** Clinical and demographic characteristics of influenza positive population.

|  | **Total JHH**<br>**population**<br>**N=94** | **Virus** |  |  |
| --- | --- | --- | --- | --- |
| **Demographics** |  | **NAg-F2H**<br>**N=34** | **NAg+F2P**<br>**N=20** | *p*-value |
| **Age**, median (range) | 54.5 (0.25-98) | 44 (2-94) | 79 (0.25-98) | 0.124 |
| **Gender female**, N (%) | 43 (46%) | 16 (47%) | 12 (60%) | 0.4083 |
| **Ethnicity, Hispanic or Latino**, N (%) | 4 (4.3%) | 3 (8.8%) | 1 (5%) | >0.99 |
| **Race**, N (%) |  |  |  | 0.63 |
| White | 43 (46%) | 13 (38%) | 11 (55%) |  |
| Black or African American | 27 (29%) | 10 (29%) | 6 (30%) |  |
| Asian | 10 (11%) | 3 (9%) | 1 (5%) |  |
| Other | 14 (15%) | 5 (15%) | 1 (5%) |  |
| **Influenza vaccination status**,<br>N/total known status (%) |  |  |  |  |
| Vaccinated | 24/33 (73%) | 4/10 (40%) | 6/7 (86%) | 0.057 |
| **Co-morbidities**, N (%) |  |  |  | 0.1 |
| None | 39 (41%) | 17 (50%) | 5 (25%) |  |
| One | 23 (24%) | 8 (24%) | 10 (50%) |  |
| Multiple | 32 (34%) | 9 (26%) | 5 (25%) |  |
| **Disease severity**, N (%) |  |  |  |  |
| Pneumonia | 14 (15%) | 3 (9%) | 4 (20%) | 0.4076 |
| Supplemental oxygen | 27 (29%) | 9 (32%) | 7 (44%) | 0.5227 |
| Admitted | 37 (39%) | 32% (11/34) | 50% (10/20) | 0.2527 |
| ICU | 9 (10%) | 3 (9%) | 2 (10%) | >0.99 |
| Hospital length of stay, median<br>(range) | 5 (1-83) | 7 (1-14) | 5 (1-13) | 0.4406 |
| Death | 5 (5%) | 1 (3%) | 2 (10%) | 0.5477 |
| Categorical variables assessed by Fisher's exact or Chi-sq. Continuous variables by Mann Whitney. Pneumonia as diagnosed by radiological findings. |  |  |  |  |

### Viruses from the 3C.2a NAg+F2P genotype associated with breathing difficulties

The clinical and demographic information from 20 patients infected with the NAg+F2P genotype was compared to the 34 patients infected with clade 3C.2a NAg-F2H genotype viruses (Table 3). The only clinical factors that differed between the patient populations was an increased reporting of wheezing and shortness of breath in individuals infected with the NAg+F2P genotype. A multivariate logistic regression analysis was performed to determine if any underlying co-morbidities were responsible for the correlations. We found no association of co-morbidities with increased wheezing or shortness of breath.

**Table 3:** Symptom comparison between individuals infected with NAg-F2H and NAg+F2P genotypes of 3C.2a H3N2 viruses.

| Symptoms, % (N reported/N total with reported) | Virus |  | *p*-value |
| --- | --- | --- | --- |
|  | NAg-F2H<br>N=34 | NAg+F2P<br>N=20 |  |
| Fever | 91% (31/34) | 71% (12/17) | 0.09 |
| Cough | 78% (25/32) | 100% (19/19) | 0.035* |
| Headache | 43% (9/21) | 40% (4/10) | >0.99 |
| Sore throat | 44% (8/18) | 38% (3/8) | >0.99 |
| Runny nose | 86% (18/21) | 73% (11/15) | 0.42 |
| Body aches | 91% (10/11) | 83% (5/6) | 1 |
| Shortness of breath | 38% (8/21) | 73% (11/15) | 0.049* |
| Wheezing | 35% (7/20) | 67% (8/12) | 0.1437 |
| Chills | 40% (8/20) | 33% (3/9) | >0.99 |
| Fatigue | 60% (12/14) | 33% (3/9) | 0.68 |
| Nausea/vomiting | 9% (2/22) | 21% (3/14) | 0.36 |
| Categorical variables assessed by Fisher's exact or Chi-sq. |  |  |  |

## Discussion

An H3N2 influenza A virus genotype, NAg+F2P, within HA clade 3C.2a with a glycosylation site on the rim of the NA sialidase pocket and the mutation H75P in its PB1-F2 protein was identified. Viruses with the NAg+F2P genotype were first detected in the 2014-15 influenza virus season and they outcompeted the NAg-F2H genotype in the 2015-16 season. This selection occurred despite NAg+F2P viruses showing reduced infectious virus production in hNEC cultures.

The main function of the influenza NA protein is to cleave sialic acid. Cleavage of sialic acid from cellular receptors leads to release of new virions and, alternatively, cleavage from decoy receptors allows virions to breach the mucosal barrier (Blumenkrantz et al., 2013; Matrosovich and Klenk, 2003). Glycosylation of NA can assist in protein folding, prevent proteolytic cleavage, mask antigenic regions, and regulate the enzyme’s catalytic activity (Chen et al., 2012; Wu et al., 2009). Antigenic escape epitopes of human seasonal N2 NA proteins have been mapped using monoclonal antibodies and molecular biology and mostly localize around the rim of the sialidase pocket (Gulati et al., 2002; Lentz et al., 1984). NA drift has also been linked to evasion of human polyclonal antibodies (Sandbulte et al., 2011). Recently, the NAg+ mutation was demonstrated to reduce NA enzymatic activity, alter virus replication fitness on hNECs, and contribute to H3N2 NA antigenic drift (Powell and Pekosz, 2020; Wan et al., 2019b). It follows that the amino acid mutations S245N and S245T, which encode a glycosylation site near the sialidase rim of the 3C.2a NA, were selected to evade human immunity despite a detrimental effect on virus replication in hNEC cultures.

PB1-F2 affects the host innate immune response as well as virus polymerase activity and transmission. The effects of PB1-F2 differ depending on protein sequence, including its length, and the cell type or animal infected. In epithelial cells, PB1-F2 can interact with mitochondrial membrane proteins ANT3 and VDAC1 and induce apoptosis by forming ion pores that release cytochrome c (Zamarin et al., 2005). PB1-F2 can disrupt MAVS signaling by interacting with MAVS, CALCOCO2, TBK1, or IRF3 thus inhibiting transcription of type I IFNs. Interactions with CALCOCO2 promote production of inflammatory cytokines through the TRAF6/NFB pathway. Alternatively, the NFkB pathway can be inhibited when PB1-F2 interacts with IKKß. PB1-F2 can also enhance viral polymerase activity. In macrophages, PB1-F2 interacts with the NLRP3 inflammasome, activates caspases, and leads to the secretion of pro-inflammatory cytokines. (Reviewed in (Kamal et al., 2018).) PB1-F2 can affect morbidity and mortality in animal models (Alymova et al., 2011). The full protein or specific residues of PB1-F2 are important for transmission in chickens, turkeys, and ferrets (Deventhiran et al., 2016; James et al., 2016; Zanin et al., 2017). Amino acid at position 75 has undergone changes in human seasonal H3N2 viruses. The 1968 pandemic H3N2 PB1 segment was of avian origin and had R75. In 1987, the PB1-F2 mutation R75H became fixed in human circulating H3N2 influenza viruses (Alymova et al., 2011). The 75^th^ AA remained H until the 2014-15 season after which, as described here, the H75P mutation became dominant. Amino acid residue 75 is located in the C-terminal portion of the PB1-F2 protein, which plays an important role in mitochondrial localization (Gibbs et al., 2003; Yamada et al., 2004) and is one of four inflammation-associated residues (Alymova et al., 2011). In the influenza virus strain A/Wuhan/359/1995, the PB1-F2 mutation H75R increased pathogenicity in mice. The lungs of mice infected with this virus also showed a trend toward reduced viral titers compared to wild type virus (Alymova et al., 2011). Here we showed that the PB1-F2 H75P mutation increased mitochondria localization.

Co-selection of the NA and PB1-F2 mutations indicates that both may contribute to virus fitness during human transmission. It is clear that the NA glycosylation allowed virus to evade antibodies at the cost of enzyme activity and reduced replication efficiency (Powell and Pekosz, 2020; Wan et al., 2019b). Since the NAg+ mutation was selected before the PB1-F2 mutation, it is worth considering if the PB1-F2 mutation was a passenger of the gene cassette or if it contributed to viral fitness. Experiments with engineered viruses that differ only by the PB1-F2 mutation H75P should be performed to determine its role in increasing transmission.

Primary cells are known to exert more biologically relevant pressures on replicating influenza viruses than the pressures exerted by transformed or immortalized cell lines. Numerous studies used primary cells to show differences in replication of engineered and isolated influenza viruses (Bui et al., 2019; Danzy et al., 2014; Elderfield et al., 2014; Fischer et al., 2015; Gerlach et al., 2017; Monteagudo et al., 2019; Wohlgemuth et al., 2017). Here, we report that hNECs were more permissive for growth of virus isolates than MDCK cell cultures. Indeed, isolation of influenza viruses in hNEC cultures allowed expansion of viruses from two thirds of positive specimens, almost three times more than the quarter that expanded after MDCK cell culture. This efficient isolation allows more virus genotypes to be studied earlier after their detection than if generating engineered viruses were necessary. We also compared the growth in hNEC cultures of two virus isolates of the NAg-F2H and NAg+F2P genotypes. This comparison showed a difference in virus fitness, but the less fit virus was the one that dominated during natural human transmission in the following season.

The 2014-15 Northern hemisphere influenza season had an unusually high number of cases and an increase in disease severity compared to the five preceding years (CDC, 2015). Overall vaccine efficacy was 19% (Zimmerman et al., 2016). This was attributed to antigenic drift of the H3N2 HA protein mediated by a new glycosylation site at amino acid 160 which partially masked neutralizing antibody epitopes (Chambers et al., 2015; Xie et al., 2015). The NAg-F2H and NAg+F2P genotypes both had the HA mutations associated with antigenic drift, but the eventual dominance of the NAg+F2P genotype suggests a selective advantage over the NAg-F2H genotype.

Associations between different human seasonal influenza A genotypes and altered disease phenotypes have been sought (Galiano et al., 2012), but rarely detected. Within human H1N1 viruses, recent pandemic strains with the HA mutation D225G associated with infection of the lower airways (Iovine et al., 2015), likely due to the increased binding of α2,3-linked sialic acid, and increased disease (van Doremalen et al., 2011). However, these viruses did not outcompete D225 encoding viruses, suggesting that the mutation did not improve transmission. The number of HA clades and frequency of isolates in each clade differ between seasons (Klein et al., 2018). The D225G and NAg+F2P associations with increased pathogenicity or symptoms appeared after a pandemic and after an antigenic drift event that each led to viral population dominance by a single genotype and its descendants. Therefore, it seems that associations between virus genotype and human disease phenotype are easier to detect and pin to specific mutations when genetic variation is small.

## Acknowledgements

We thank Samantha Lycett (University of Edinburgh, UK) and Colin Russell (Cambridge University, UK) for sharing their phylogenetic tree branch coloring protocols as well as Monica Galiano (Public Health England) for the discussion about correlations between genotype and phenotype. We are grateful to Anne Hamacher-Brady and Nathan Brady for assistance with microscopy and the members of the Pekosz laboratory, Sabra Klein Laboratory and Kimberley Davis laboratory for helpful discussions. This work was supported by the NIH/NIAID Center of Excellence in Influenza Research and Surveillance contract HHS N272201400007C (Johns Hopkins University) and the Richard Eliasberg Family Foundation.

